# Divergence of Cortical Force-Generating Mechanisms Underlies Differences in Spindle Behavior between *C. elegans* and *C. inopinata*

**DOI:** 10.64898/2026.06.06.730595

**Authors:** Shun Oomura, Koji Kyoda, Shuichi Onami, Nami Haruta, Asako Sugimoto

## Abstract

Microtubule-dependent pronuclear migration and mitotic spindle positioning are fundamental processes during the first embryonic division in many animals. In the one-cell embryo of *Caenorhabditis elegans*, these events are regulated by well-characterized pulling forces acting on astral microtubules, including cortical forces mediated by the Gα-GPR-LIN-5 dynein complex. Although the overall framework of these dynamics is conserved, recent studies have revealed substantial interspecies variation in their regulation. Here, we investigated nuclei and mitotic spindle behaviors in one-cell embryos of *Caenorhabditis inopinata*, the closest known relative of *C. elegans*, using live-cell imaging and functional perturbation. We found that *C. inopinata* embryos exhibit altered pronuclear migration, reduced anaphase spindle oscillations, and slower centrosome diffusion during telophase compared with *C. elegans*. These differences suggest weaker cortical pulling forces. Functional analyses using RNA interference showed that GPR retains its essential role in force generation, whereas the contribution of the microtubule depolymerizing kinesin KLP-7 is reduced in *C. inopinata*. Our results point to evolutionary changes in microtubule-regulated spindle dynamics, and provide insight into how conserved cellular processes can diversify through subtle changes in their underlying mechanisms.

## Introduction

Precise regulation of microtubules is essential for the spatiotemporal control of intracellular dynamics. These structures play a central role in key events during early embryogenesis, including pronuclear migration and mitotic spindle positioning.

In the one-cell-stage embryo of *Caenorhabditis elegans*, nuclear and mitotic spindle dynamics are regulated by a combination of cytoplasmic and cortical forces acting on astral microtubules. Upon fertilization, centrosomes near the male pronucleus nucleate microtubules that guide the female pronucleus toward the posterior, resulting in the formation of a nucleus-centrosome complex (NCC) [1]. This complex then migrates to the cell center and rotates along the anterior-posterior axis. The spindle is positioned by two distinct types of forces: (1) cytoplasmic, length-dependent forces involving cytoplasmic dynein along microtubules [2], and (2) cortical pulling forces generated by interactions between astral microtubules and the Gα-GPR-LIN-5 dynein complex at the cell cortex [3]. Posterior enrichment of this complex causes net posterior displacement of the spindle and generates transverse oscillations during anaphase [3-8].

Although the molecular components of this force-generating machinery are broadly conserved, studies across Rhabditida nematodes have revealed substantial variation in spindle dynamics, suggesting evolutionary flexibility in the regulation of these forces [9-11]. Understanding the basis of this variation provides an opportunity to investigate how microtubule regulatory mechanisms have diversified across species.

*Caenorhabditis inopinata*, the closest known relative of *C. elegans*, provides a useful model for such comparative studies. Despite their close phylogenetic relationship, having diverged ∼10.5 million years ago, these two species differ in adult morphology, reproductive systems, and ecological adaptations [12]. These differences suggest possible divergence in the intracellular dynamics, including microtubule-dependent events. However, the intracellular dynamics during *C. inopinata* development remains poorly characterized.

In this study, we conducted a comparative analysis of the dynamics of pronuclear migration and mitotic spindles in one-cell-stage embryos of *C. inopinata* and *C. elegans* using differential interference contrast (DIC) and fluorescence live imaging. We identified differences in pronuclear migration and mitotic spindle behavior, suggesting that cortical pulling forces are weaker in *C. inopinata*. RNAi-based assays further suggested that reduced activity of the microtubule-depolymerizing kinesin KLP-7 may contribute to this weakened pulling force. These findings demonstrate that even subtle changes in microtubule regulatory pathways can lead to divergent intracellular behaviors between closely related species.

## Materials and methods

### Strains and culture conditions

*C. elegans* strains were cultured as described by Brenner (1974) [13]. Worms were cultured at 20 °C on nematode growth medium (NGM) plates seeded with *Escherichia coli* OP50. The following strains were used: N2 (wild-type), SA1161 (*tjSi193[eft-3p::gfp::his-58::tbb-2 3’UTR; rps-27p::NeoR::unc-54 3’UTR]*) [14], and SA250 (*tjIs54[pie-1p::gfp::tbb-2; pie-1p::2xmCherry::tbg-1; unc-119+]; tjIs57[pie-1p::mCherry::his-48; unc-119+]*) [15]. The wild-type strain of *C. briggsae* (HK104) was cultured under the same conditions.

*C. inopinata* strains were cultured at 27 °C on modified NGM plates, as described by Oomura et al. (2022) [14], seeded with *E. coli* HT115 (DE3). The following strains were used: NKZ35 (wild-type), SA1438 (*tjIs372[eft-3p::gfp::his-58::tbb-2 3’UTR; rps-0p::HygR::unc-54 3’UTR]*) [14], and SA1648 (*tjIs375[pie-1p::gfp::tbb-2::tbb-2 3’UTR; rps-0p::HygR::unc-54 3’UTR]*) [14].

### Microscopy

For live-cell imaging, gravid *C. elegans* hermaphrodites or gravid *C. inopinata* females were dissected in M9 buffer to release embryos, which were then mounted on 2% agarose pads. DIC images were acquired every 0.5 s using an ORCA-R2 digital CCD camera (Hamamatsu Photonics) mounted on an Axioplan 2 imaging microscope (Zeiss) equipped with a C-Apochromat 63x/1.2 W Corr M27 water-immersion objective lens (Zeiss).

Fluorescent live images of GFP::histone and GFP::*β*-tubulin were acquired using a CSU-X1 spinning disc confocal system (Yokogawa Electric Corporation). GFP::histone images were acquired every 30 s with 23 z-slices at 1 μm intervals using an iXonEM+ EMCCD camera (Andor technology) mounted on an Axioplan 2 imaging microscope (Zeiss) equipped with a C-Apochromat 63x/1.2 W Corr M27 water-immersion objective lens (Zeiss). GFP::*β*-tubulin images were acquired every 30 s for non-RNAi-treated *C. elegans* embryos or 60 s for RNAi-treated *C. elegans* and *C. inopinata* embryos with 10 z-slices at 1 μm intervals using an ORCA-flash4.0 V3 digital CMOS camera (Hamamatsu Photonics) mounted on an IX71 microscope (Olympus) equipped with a UPlanSApo 60×/1.30 silicone oil-immersion objective lens (Olympus). MetaMorph software (Molecular Devices) was used to control the microscopes. All live imaging experiments were performed at 25 °C unless otherwise noted.

### Tracking of pronuclei, chromosomes, centrosomes, and spindle movements

ImageJ/Fiji software (NIH; https://imagej.net/Fiji) was used for image processing and analysis. Fluorescent images were processed by z-projection using averaged intensity values across z-slices.

Pronuclei, centrosomes, and chromosomes were manually tracked, and their positions were recorded relative to embryo length (anterior-posterior axis) and width (dorsal-ventral axis). Spindle position was defined as the midpoint between the two centrosomes, spindle angle as the angle of the line connecting the centrosomes, and spindle length as the distance between the centrosomes.

### Quantification of mitotic spindle oscillation

Centrosome positions were tracked at 1-s intervals in DIC images from nuclear envelope break down (NEBD) to the completion of cytokinesis. In *C. inopinata*, rocking peaks were defined as the time points at which a continuous increase (or decrease) in spindle angle lasting at least 4 s switched to a continuous decrease (or increase) lasting at least 4 s. The maximum angular displacement was defined as the largest change in spindle angle between consecutive rocking peaks. Oscillation frequency was determined as the dominant frequency component between 0.04 and 0.10 Hz by Fast Fourier transform (FFT) with Hamming window correction.

### Statistical analysis

Quantification method, sample sizes, and statistical analyses are described in the main text and figure legends. Data were analyzed using GraphPad Prism 9. Statistical significance was assessed using one-way ANOVA followed by Tukey’s multiple comparisons test.

### Ortholog search

Orthologs of *C. elegans* genes in the *C. inopinata* genome were identified using BLAST+ search [16-18] with amino acid sequences from the *C. inopinata* v7.10 gene set [12]. The amino acid sequence of the longest isoform of each *C. elegans* gene was used as the query, and the *C. inopinata* gene with the lowest E-value was selected as the ortholog. *C. inopinata* orthologs are referred to with the prefix “*Cin*”.

### RNA interference

For *C. elegans*, RNAi was performed using the soaking method [19]. Target sequences of *gpr-1* (1063 bp), *let-99* (1024 bp), and *klp-7* (1025 bp) were amplified from cDNA. dsRNA was synthesized using T7 RiboMAX Express Large Scale RNA Production System (Promega) and purified by phenol-chloroform extraction. Twenty L4 hermaphrodites were soaked in RNAi solution containing 2 µg/µl dsRNA at 24.5 °C for 24 h, recovered, and then incubated on NGM plates at 24.5 °C for an additional 24 h before imaging. To assess embryonic lethality, five adult hermaphrodites were allowed to lay eggs at 24.5 °C for 2 h, and the unhatched rate was measured 24 h later.

For *C. inopinata*, RNAi was performed using the feeding method [12, 20], because the recovery rate was very low for the soaking method. Target sequences of *Cin-gpr-1* (1017 bp), *Cin-let-99* (1040 bp), and *Cin-klp-7* (1040 bp) were amplified from genomic DNA and cloned into the T444T feeding RNAi vector [21]. *E. coli* HT115(DE3) transformed with each RNAi plasmid was seeded onto RNAi plates (NGM with 1 mM IPTG) and incubated at room temperature for one day. Twenty gravid females were transferred to the RNAi plates, and their progeny were cultured at 27 °C for 96-120 h until they reached the gravid adult stage for imaging. To assess embryonic lethality, 20 females and 10 males were mated and allowed to lay eggs at 27 °C for 8 h, and the unhatched rate was measured 24 h later.

## Results

### The asymmetry of the first embryonic division in *C. inopinata* is similar to that in *C. elegans*

We first compared egg size, the duration of the first cell division, and division asymmetry between one-cell-stage embryos of *C. elegans* (N2) and *C. inopinata* (NKZ35) using differential interference contrast (DIC) microscopy (Fig 1). Although *C. inopinata* adults are more than twice as long as *C. elegans* adults [12], the egg length of *C. inopinata* (58.8 ± 3.6 µm, mean ± SD, *n* = 11) was only 5.7% longer than that of *C. elegans* (55.7 ± 2.9 µm, *n* = 10, *p* < 0.05), and egg width did not differ significantly between the two species (*C. elegans*: 32.5 ± 1.3 µm, *n* = 10; *C. inopinata*: 32.2 ± 0.8 µm, *n* = 11) (Fig 1B). Because optimal growth temperatures differ between the two species (*C. elegans*: 15-24°C; *C. inopinata*: 25-29°C), all observations were conducted at 25°C. Under these conditions, the duration of the first cell division, measured from nuclear envelope breakdown to cytokinesis, was 1.6-fold longer in *C. inopinata* (7.7 ± 0.4 min, *n* = 9) than in *C. elegans* (4.7 ± 0.4 min, *n* = 10, *p* < 0.0005) (Fig 1C). The position of the first cleavage furrow, expressed as the relative position (%) = distance from the anterior end of the embryo / embryo length, was 56.0 ± 0.9% (*n* = 11) in *C. inopinata* and 56.7 ± 1.4% (*n* = 10) in *C. elegans*, indicating no significant difference in the asymmetry of the first embryonic division between the two species (Fig 1D).

**Fig 1.**
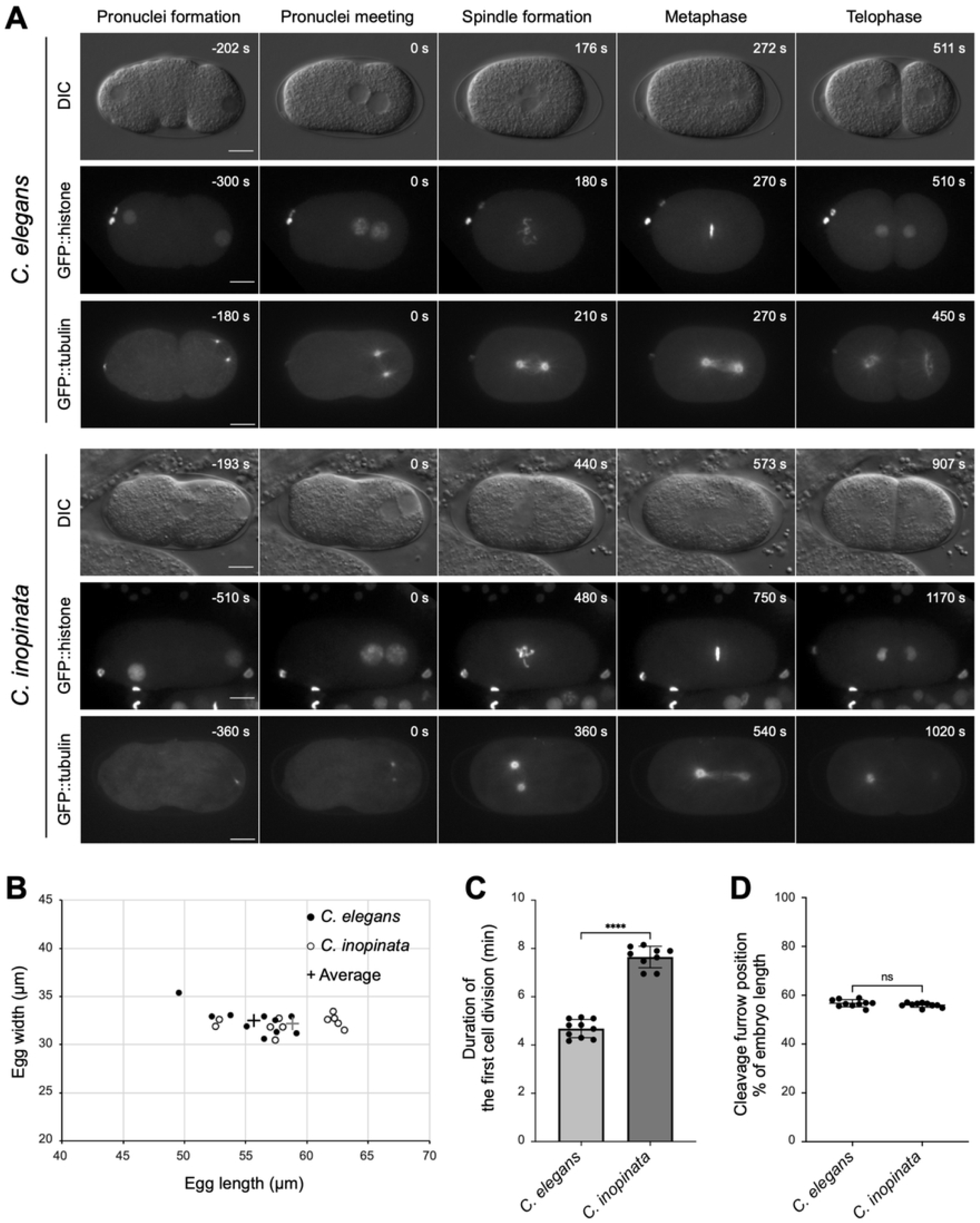
The first asymmetric cell division in *C. elegans* and *C. inopinata* embryos. (A) DIC live imaging of wild-type embryos and fluorescent live imaging of embryos expressing GFP::histone or GFP::*β*-tubulin. Embryos are oriented with the anterior side to the left. Time is shown relative to the pronucleus meeting (s). All images were acquired at 25°C. Scale bars, 10 µm. (B) Egg length and width measured from DIC images of *C. elegans* (*n* = 10) and *C. inopinata* (*n* = 11) embryos. (C) Duration of the first cell division, defined as the time from NEBD to the completion of cytokinesis, based on DIC live imaging of *C. elegans* (*n* = 10) and *C. inopinata* (n = 9) embryos. ****P < 0.001; unpaired *t*-test. (D) Cleavage furrow position measured from DIC images of *C. elegans* (*n* = 10) and *C. inopinata* (*n* = 9) embryos. ns, no significant; unpaired *t*-test.

### Pronucleus dynamics differ between *C. elegans* and *C. inopinata* embryos

Next, we compared pronuclear dynamics between one-cell-stage embryos of *C. elegans* and *C. inopinata* using the fluorescent live-cell imaging of GFP::histone (Fig 1A, Fig 2A). In *C. elegans*, the female and male pronuclei formed at the anterior and posterior poles, respectively (Fig 2A). In *C. inopinata*, the male pronucleus formed at the posterior pole, as in *C. elegans*, whereas the female pronucleus formed at various positions; in more than half of the embryos, it formed on the posterior side (6/10) (Fig 2A). The position of the female pronucleus in one-cell-stage embryos reflects the position of the nucleus in mature oocytes. In *C. elegans*, the oocyte nucleus migrates from the center to the distal cortex during oocyte maturation (Fig 2C, D). In contrast, in *C. inopinata*, the oocyte nucleus did not migrate distally, even in oocytes adjacent to the spermatheca (Fig 2C, D). This difference may explain the various positions of female pronucleus formation. In both species, the female and male pronuclei move toward each other at the posterior side. However, the male pronucleus exhibited less anterior movement in *C. inopinata* than in *C. elegans* (Fig 2A). Consequently, the pronuclei met at a more posterior in *C. inopinata* (80.1 ± 3.8%, *n* = 10) than in *C. elegans* (66.2 ± 3.4%, *n* = 10; *p* < 0.0005) (Fig 2B).

**Fig 2.**
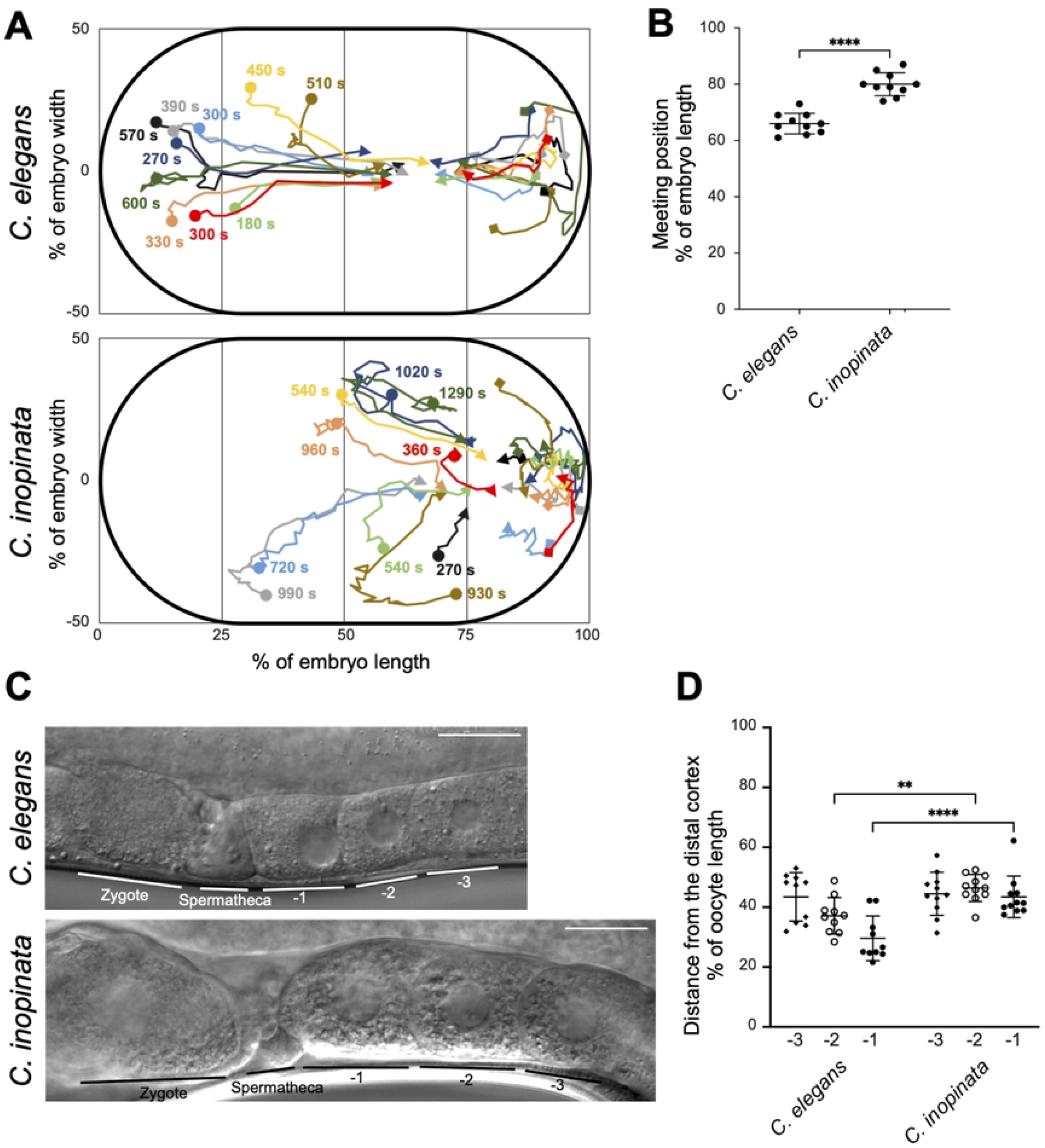
Pronucleus dynamics in *C. elegans* and *C. inopinata* embryos. (A) Time course of pronucleus movement. Pronucleus positions were tracked every 30 s using fluorescent live imaging of embryos expressing GFP::histone: *C. elegans* (*n* = 10), *C. inopinata*, (*n* = 10). The horizontal axis indicates the relative position (%) along the embryo length, and the transverse axis indicates the relative position (%) along the embryo width. Symbols indicate the following positions: circle, female pronucleus formation; square, male pronucleus formation; arrowhead, pronucleus meeting. Each colored line represents an individual embryo. The number near the female pronucleus formation indicates the time in seconds until the pronucleus meeting. (B) Position of pronucleus meeting: *C. elegans* (*n* = 10), *C. inopinata*, (*n* = 10). ****P < 0.001; unpaired *t*-test. (C) DIC images of the proximal gonad in adult *C. elegans* hermaphrodites and *C. inopinata* females. -1, - 2, and -3 indicate proximal oocytes adjacent to the spermatheca. Scale bars, 100 µm. (D) Distance between the oocyte nucleus and the distal cortex in proximal oocytes: *C. elegans* (*n* = 10), *C. inopinata* (*n* =11). Oocytes at the corresponding positions were compared between *C. elegans* and *C. inopinata*. **P < 0.01; ****P < 0.0001; unpaired *t*-test.

### NCC and mitotic spindle movements differ between *C. elegans* and *C. inopinata* embryos

In *C. elegans*, after pronucleus meeting, the nucleus-centrosome complex (NCC) migrates toward the center of the embryo, a process known as centration, and rotates 90 degrees to align the centrosomes along the anterior-posterior (A/P) axis. The NCC then undergoes nuclear envelope breakdown (NEBD), and the mitotic spindle forms along the A/P axis. The spindle is displaced slightly posteriorly by the end of anaphase. To compare NCC and spindle movements between *C. elegans* and *C. inopinata* embryos, we performed fluorescent live-cell imaging of GFP:: *β*-tubulin, as a microtubule marker (Fig 1A, Fig 3A). During centration, the NCC in *C. inopinata* migrated anteriorly by 39.9 ± 6.3% of the embryo length (*n* = 7), approximately twice the distance observed in *C. elegans* (20.3 ± 1.9%, *n* = 9), and reached the anterior side of the embryo (Fig 3A). Consequently, the spindle formed more anteriorly in *C. inopinata* (42.1 ± 5.2%, *n* = 11) than in *C. elegans* (52.1 ± 1.5%, *n* = 10, *p* < 0.0005), a phenotype referred to here as “overcentration” (Fig 3B). In *C. elegans*, the NCC rotated to align the centrosomes along the A/P axis (initial spindle angle; 9.1 ± 6.2 degrees, *n* = 10). In contrast, the NCC in *C. inopinata* exhibited limited rotation, resulting in a larger initial spindle angle (54.3 ± 25.4 degrees, *n* = 10, *p* < 0.0005) (Fig 3B). The initial spindle length did not differ significantly between the species (*C. elegans*: 10.0 ± 0.7 µm, *n* = 10; *C. inopinata*: 10.5 ± 1.0 µm, *n* = 11) (Fig 3B). By the end of metaphase, however, the spindle became longer in *C. inopinata* (17.1 ± 1.3 µm, *n* = 10) than in *C. elegans* (12.1 ± 0.8 µm, *n* = 10, *p* < 0.0005) (Fig 3C). At this stage, the spindle in *C. inopinata* was positioned slightly posterior to the center of the embryo (58.9 ± 2.3%, *n* = 11), similar to its position in *C. elegans* (57.1 ± 1.8%, *n* = 10), and was aligned along the A/P axis (Fig 1A, Fig 3D).

**Fig 3.**
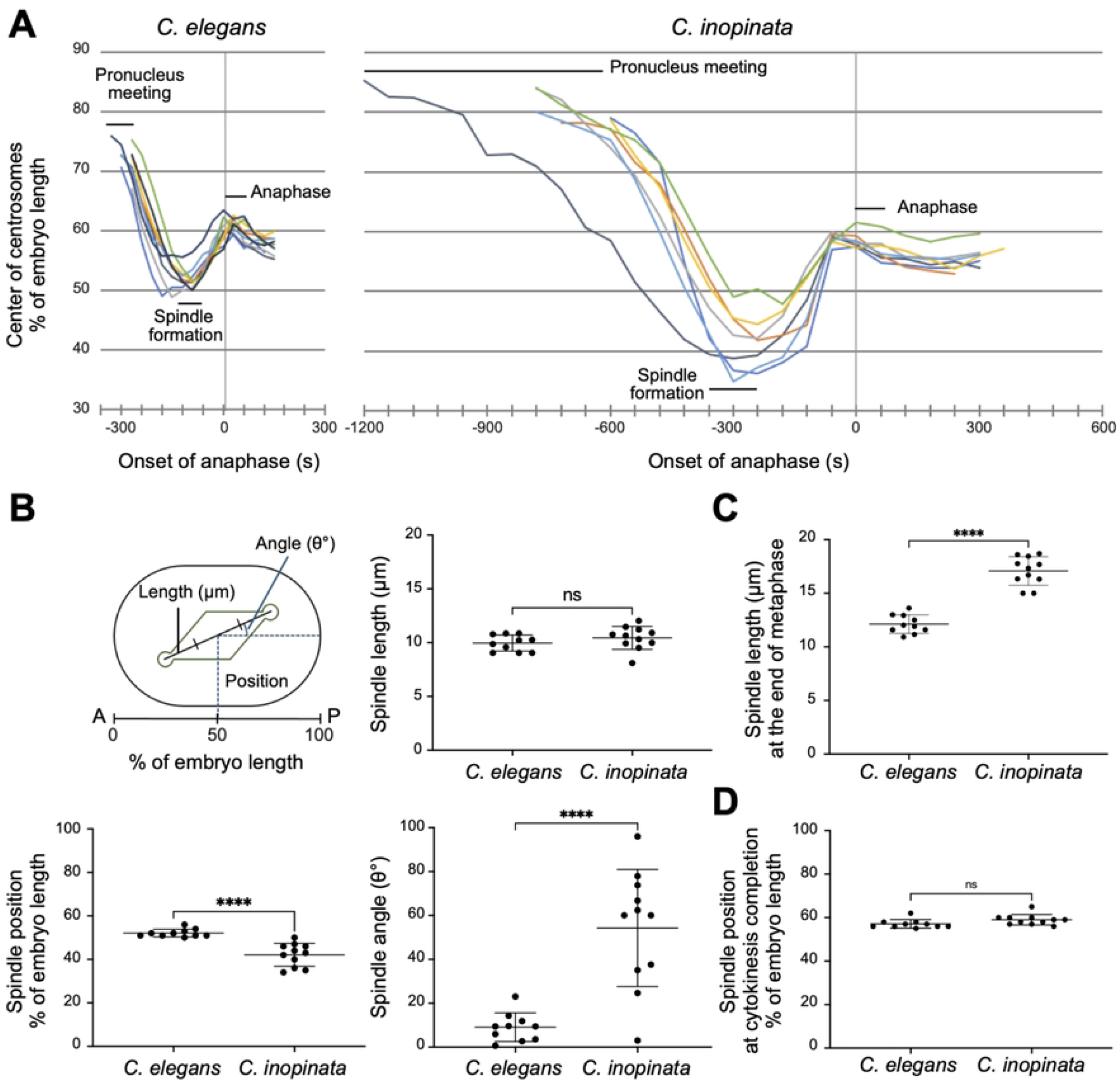
NCC and spindle dynamics in *C. elegans* and *C. inopinata* embryos. (A) Time course of spindle positions from pronucleus meeting to the completion of cytokinesis. Spindle positions were tracked using fluorescent live imaging of GFP::*β*-tubulin every 30 s in *C. elegans* (*n* = 9) and 60 s in *C. inopinata* (*n* = 7). Spindle positions are shown as relative positions (%) along the embryo length. Time is indicated relative to anaphase onset (t = 0 s). (B) Spindle position, angle, and length at spindle formation. The measurement model is shown in the upper left panel. *C. elegans* (*n* = 10), *C. inopinata* (*n* = 11). ****P < 0.0001; ns, not significant; unpaired *t*-test. (C) Spindle length at the end of metaphase. *C. elegans* (*n* = 10), *C. inopinata* (*n* = 11). ****P < 0.0001; unpaired *t*-test. (D) Spindle position at the completion of cytokinesis. *C. elegans* (*n* = 10), *C. inopinata* (*n* = 11). ns, not significant; unpaired *t*-test.

### Mitotic spindle oscillation is smaller in *C. inopinata* than in *C. elegans* and *C. briggsae*

During anaphase, cortical microtubule pulling forces induce transverse spindle oscillations in *C. elegans* [4, 8]. To compare spindle oscillation among species, we tracked centrosomes using DIC live imaging in *C. inopinata, C. elegans*, and *C. briggsae*, the latter of which is known to show smaller oscillations than *C. elegans* [11, 22] (Fig 4A). The number of rocking cycles in *C. inopinata* (3.4 ± 1.5, *n* = 9) was similar to that in *C. briggsae* (3.7 ± 0.8, *n* = 9) but significantly lower than in *C. elegans* (5.8 ± 1.4, *n* = 10, *p* < 0.005) (Fig 4B). The maximum angular displacement in *C. inopinata* (13.1 ± 5.1°, *n* = 9) was similar to that in *C. briggsae* (14.7 ± 7.1°, *n* = 9) but significantly smaller than that in *C. elegans* (26.8 ± 11.7°, *n* = 10, *p* < 0.005) (Fig. 4B). The oscillation frequency in *C. inopinata* (38.7 ± 6.0 mHz, *n* = 9) was significantly lower than those in both *C. elegans* (56.8 ± 7.1 mHz, *n* = 10, *p* < 0.0005) and *C. briggsae* (62.2 ± 4.3 mHz, *n* = 9, *p* < 0.0005) (Fig 4B). Taken together, the spindle oscillation in *C. inopinata* is smaller than in *C. elegans*, possibly reflecting differences in the interaction between astral microtubules and the cell cortex between the two species.

**Fig 4.**
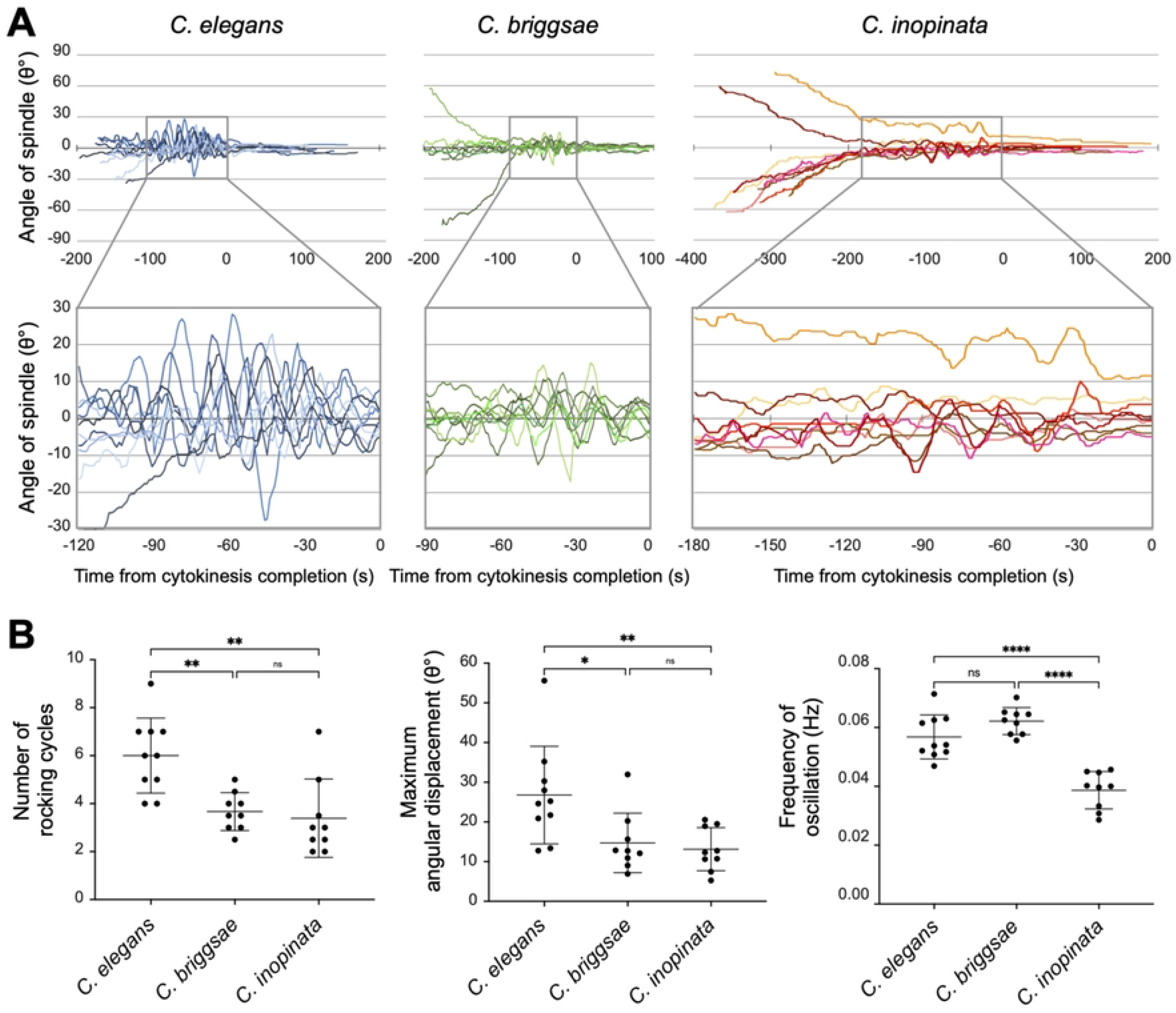
Comparison of mitotic spindle oscillation parameters in *C. elegans, C. inopinata*, and *C. briggsae* embryos. (A) Time course of spindle angle. The upper panel shows the period from NEBD to the completion of cytokinesis. The lower panel focuses on the oscillation period. Time is indicated relative to the completion of oscillation (t = 0 s). The spindle angle was calculated every second using DIC live imaging. *C. elegans* (*n* = 10), *C. inopinata* (*n* = 9), *C. briggsae* (*n* = 9). (B) Comparison of the number of rocking cycles, maximum angular displacement (θ°), and oscillation frequency (Hz). Frequency was determined by fast Fourier transform. *C. elegans* (*n* = 10), *C. inopinata* (*n* = 9), *C. briggsae* (*n* = 9). *P < 0.05; **P < 0.01; ****P < 0.0001; ns, not significant; Tukey’s multiple comparisons test.

### Centrosome disassembly during telophase occurs more slowly in *C. inopinata* than in *C. elegans*

We next compared centrosome disassembly during telophase, which is thought to be associated with the strength of cortical microtubule pulling forces [23, 24], using fluorescent live-cell imaging of GFP::*β*-tubulin. In *C. elegans*, the posterior bias of cortical pulling forces led to the rapid expansion and fragmentation of the posterior centrosome, followed by the anterior centrosome (*n* = 10) (Fig 5). In *C. inopinata*, the posterior centrosome also disassembled before the anterior centrosome, but both centrosomes diffused slowly without marked expansion or fragmentation (*n* = 11) (Fig 5). These results suggest that cortical pulling forces in *C. inopinata* are biased posteriorly, as in *C. elegans*, but are weaker overall.

**Fig 5.**
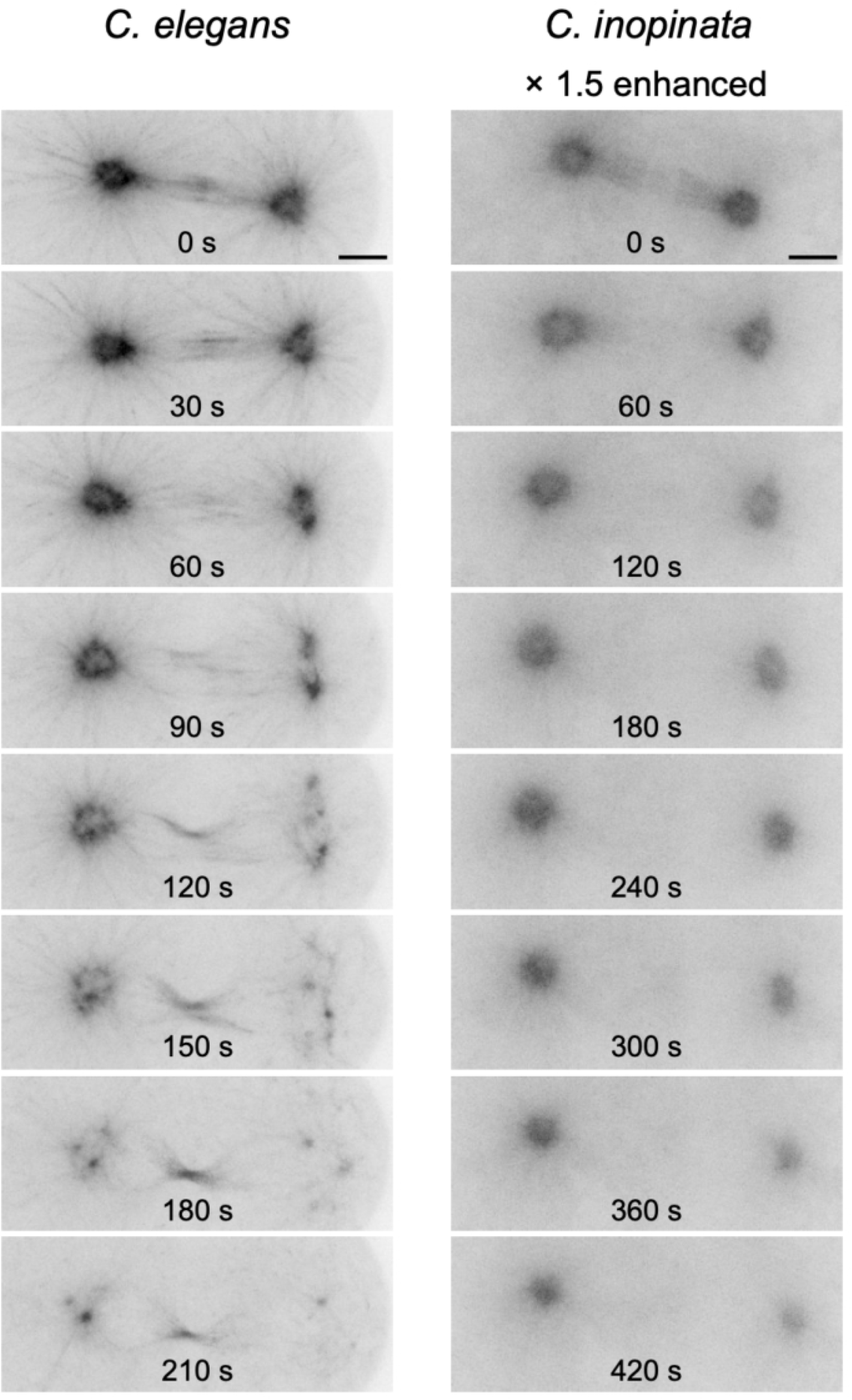
Centrosome disassembly during telophase in *C. elegans* and *C. inopinata* embryos. Centrosomes were recorded using fluorescent live imaging of embryos expressing GFP::*β*-tubulin every 30 s in *C. elegans* and 60 s in *C. inopinata*. Time is indicated relative to anaphase onset (s). Scale bars, 5 µm.

### Comparative analysis of genes involved in the regulation of cortical microtubule pulling forces

The smaller mitotic spindle oscillation during anaphase and the slower posterior centrosome dispersal were consistent with weaker cortical microtubule pulling forces in *C. inopinata* than in *C. elegans*. In *C. elegans*, the Gα-GPR-LIN-5 complex functions as a cortical pulling force generator, and its posterior enrichment is regulated by multiple protein interactions. In addition, spindle oscillation is affected by microtubule dynamics [25-28]. Genes involved in these processes were generally conserved between *C. elegans* and *C. inopinata* (Table 1). We therefore hypothesized that the contribution of each gene might differ between the species. To test this hypothesis, we selected three genes, *gpr-1/2, let-99*, and *klp-7*, and compared their reduction-of-function phenotypes by RNAi. In *C. elegans, gpr-1/2* encode two functionally redundant GPR proteins [3], whereas *C. inopinata* has a single ortholog. LET-99 inhibits both GPR and LIN-5 localization [3, 5]. KLP-7 is a kinesin-13 microtubule depolymerization factor involved in regulating astral microtubule levels [27, 28].

**Table 1:**
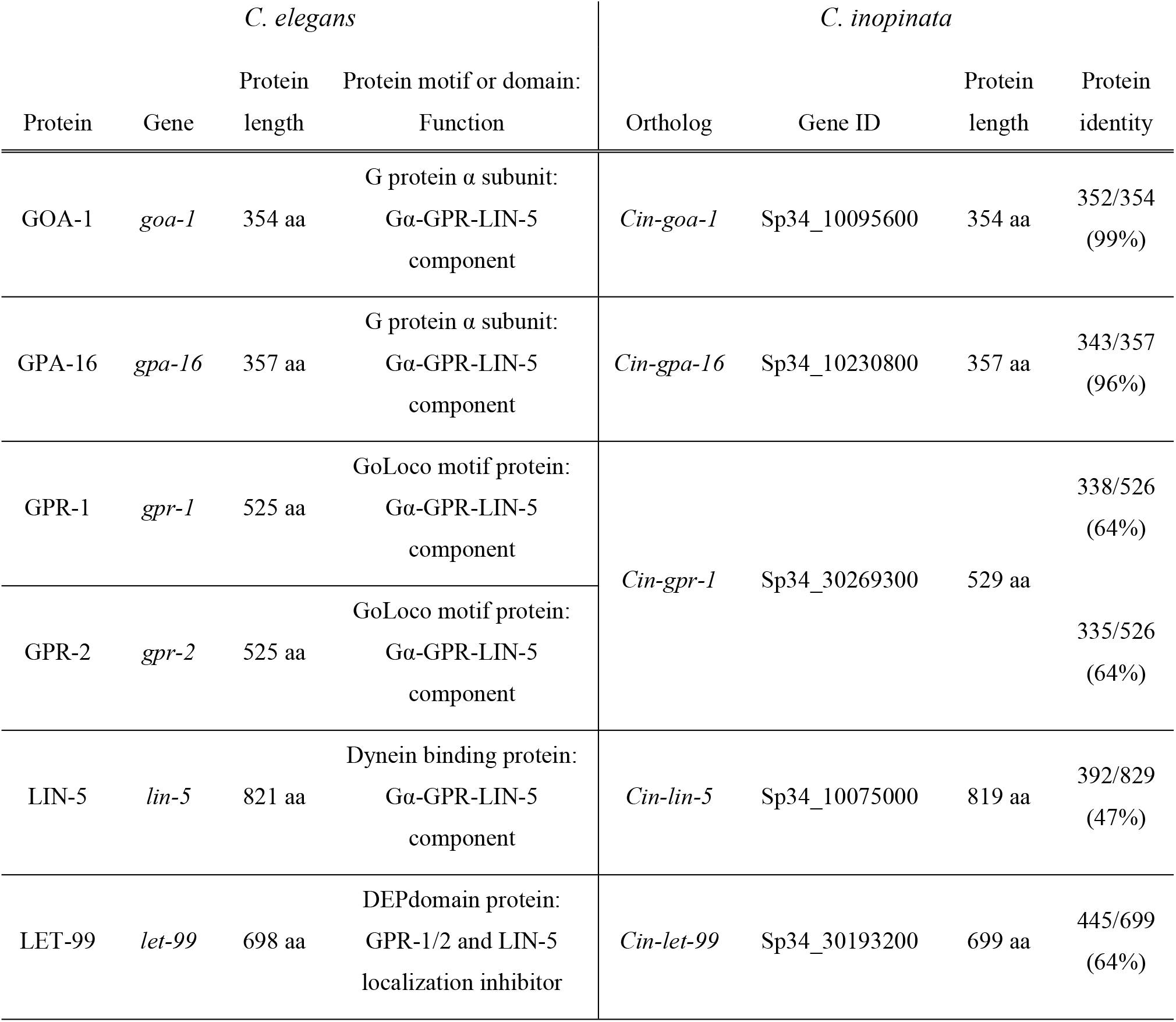
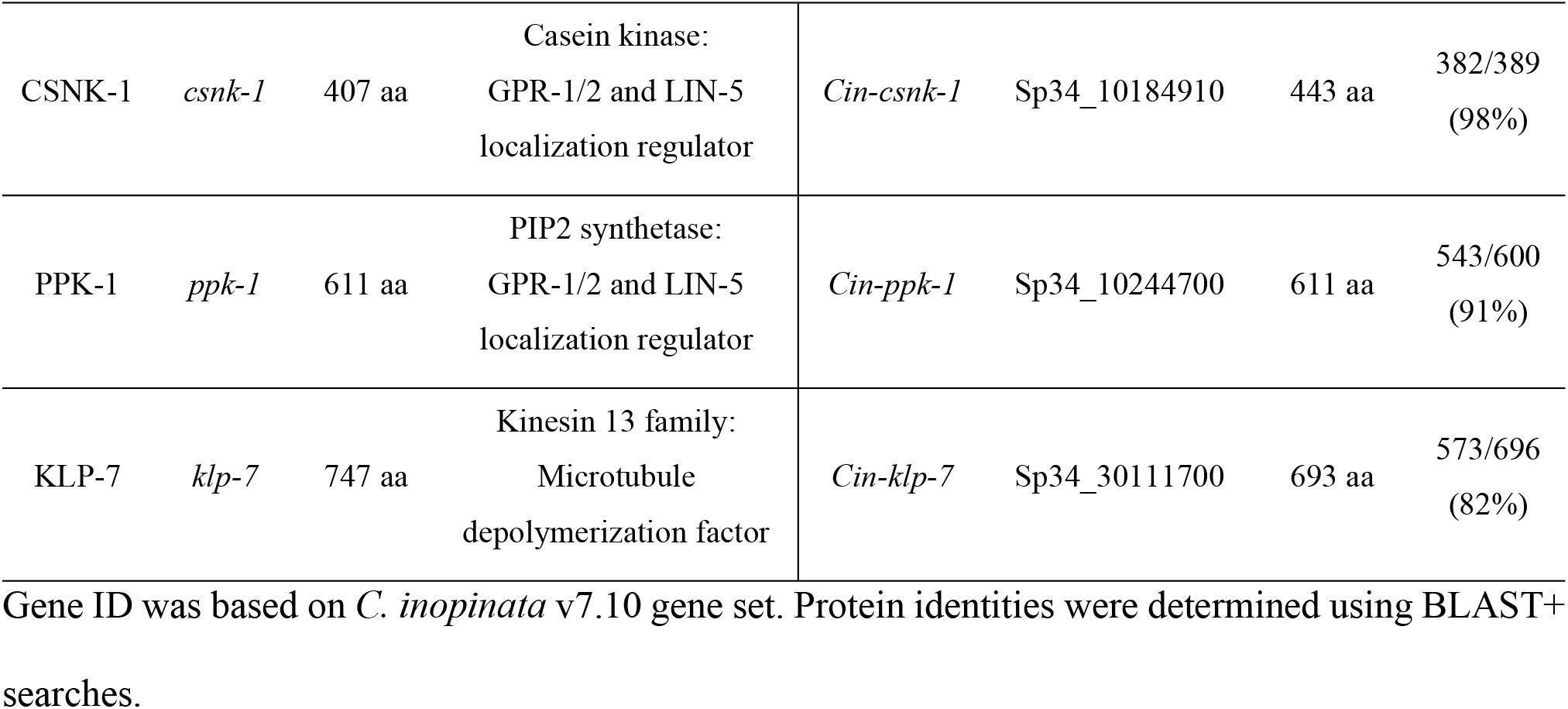
Genes involved in the control of cortical microtubule pulling forces in *C. elegans*.

RNAi knockdown of *Cin-gpr-1, Cin-let-99*, and *Cin-klp-7* resulted in high embryonic lethality in *C. inopinata*, similar to the phenotypes observed for their orthologs in *C. elegans* (Fig 6A). In *C. elegans, gpr-1(RNAi)* embryos exhibited reduced spindle oscillations and posterior spindle displacement, whereas *let-99(RNAi)* embryos exhibited earlier onset and increased amplitude of spindle oscillations. Similar phenotypes were observed in *Cin-gpr-1(RNAi)* and *Cin-let-99(RNAi)* embryos, suggesting that the role of *Cin-gpr-1* and *Cin-let-99* are conserved in *C. inopinata*. In contrast, although *klp-7(RNAi)* embryos of *C. elegans* exhibited spindle misalignment along the A/P axis and most failed to complete cell division (6/8 embryos), *Cin-klp-7(RNAi)* embryos of *C. inopinata* completed asymmetric cell division without significant defects in spindle dynamics (Fig 6B). These results suggest that although *Cin-klp-7* is essential for embryogenesis in *C. inopinata*, its contribution to the first asymmetric division is limited.

**Fig 6.**
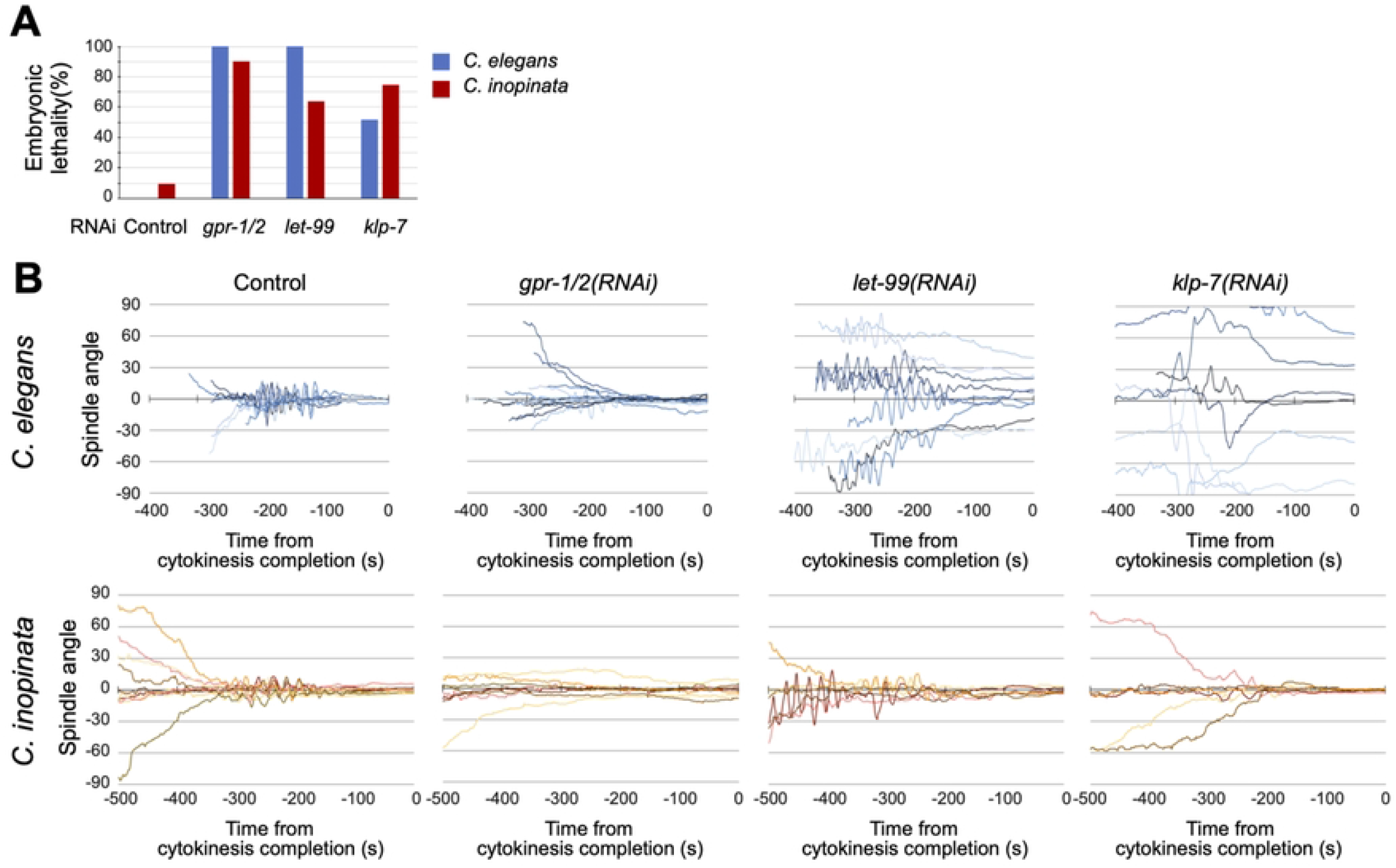
Reduction-of-function phenotypes of *gpr-1/2, let-99*, and *klp-7*, and their orthologs in *C. elegans* and *C. inopinata* embryos. (A)Embryonic lethality observed in each RNAi experiment. For *C. elegans* soaking RNAi, each value represents the mean of four independent measurements: Control, 0% (0/47, 0/43, 0/50, 0/37); *gpr-1/2(RNAi)*, 100% (48/48, 26/26, 48/48, 32/32); *let-99(RNAi)*, 100% (29/29, 38/38, 33/33, 45/45); *klp-7(RNAi)*, 51.5% (22/46, 31/48, 22/52, 9/17). For *C. inopinata* feeding RNAi, each value represents the mean of two independent measurements: control, 9.5% (4/97, 19/127); *Cin-gpr-1(RNAi)*, 90.1% (25/28, 30/33); *Cin-let-99(RNAi)*, 63.9% (75/125, 84/124); *Cin-klp-7(RNAi)*, 74.7% (72/103, 70/88). (B)Time course of spindle angle in each RNAi experiment. The upper panel shows *C. elegans* from NEBD to cytokinesis completion (t = 0 s). The lower panel shows *C. inopinata* during the 500 s preceding cytokinesis completion (t = 0 s). The spindle angle was measured every second using DIC live imaging. *C. elegans*: control, n = 16; *gpr-1/2(RNAi), n* = 13; *let-99(RNAi), n* = 9; *klp-7(RNAi), n* = 8. *C. inopinata*: control, *n* = 10; *Cin-gpr-1(RNAi), n* = 8; *Cin-let-99(RNAi), n* = 6; *Cin-klp-7(RNAi), n* = 6.

## Discussion

In this study, we identified differences in microtubule-dependent pronuclear migration and mitotic spindle dynamics in one-cell-stage embryos of *C. elegans* and *C. inopinata*. In *C. inopinata*, the female pronucleus formed at various positions and met the male pronucleus at a more posterior position than in *C. elegans*. After pronucleus meeting, the NCC migrated toward the anterior side of the embryo and showed limited rotation, resulting in spindle formation at a more anterior position than in *C. elegans*. During cell division, spindle oscillations during anaphase were smaller, and centrosome dispersal during telophase occurred more slowly in *C. inopinata* than in *C. elegans*. These observations are consistent with the possibility that cortical microtubule pulling forces are weaker in *C. inopinata* than in *C. elegans*. RNAi-based assays further suggested that reduced activity of the microtubule-depolymerizing kinesin KLP-7 in *C. inopinata* may contribute to this difference in microtubule pulling force.

In *C. elegans*, spindle oscillation is caused by cortical microtubule pulling forces generated by the Gα-GPR-LIN-5 complex. Our RNAi experiments indicated that, in *C. inopinata*, this complex is also involved in spindle positioning, but its force-generating activity may be reduced. This reduction may result from a lower force-generating capacity of the Gα-GPR-LIN-5 complex in *C. inopinata*, or from fewer centrosomal microtubules interacting with the complex. We also found that the microtubule depolymerization factor KLP-7 contributes less to positioning of the first mitotic spindle in *C. inopinat*a. This finding suggests that the regulation of centrosomal microtubules differs between the two species, which may affect the interaction between centrosomal microtubules and cortical force generators.

In addition to differences in spindle dynamics, we found overcentration of the NCC in *C. inopinata*. In *C. elegans*, anterior movement of the NCC is mainly driven by microtubule length-dependent pulling forces [2]. It is therefore possible that the regulation of length-dependent forces differs between these species. Further RNAi experiments targeting factors involved in microtubule length-dependent pulling forces, such as cytoplasmic dyneins and kinesins, will help clarify the molecular basis of these differences.

Comparative analysis among 34 Rhabditid species related to *C. elegans* has shown diverse cellular behaviors during early cell division. However, two features, 90-degree NCC rotation (31/34 species) and NEBD at the center of the embryo (30/34), are well conserved [10]. Notably, these features were rarely observed in *C. inopinata*. Among *Caenorhabditis* nematodes, *C. inopinata* exhibited one of the most anterior spindle formation positions and one of the smallest spindle oscillations [11]. Thus, the cellular behaviors of *C. inopinata* during the first cell division have diverged substantially from those of closely related species.

In *C. inopinata*, transposable elements are desilenced and expanded [12, 29, 30], which is believed to contribute to faster phenotype evolution than other related species. For example, optimal growth temperature of *C. inopinata* is 25-29°C, which is significantly higher than that of *C. elegans* (15-24°C) [12]. Because microtubule dynamics are temperature-sensitive, adaptation to higher temperatures and the regulatory mechanisms of microtubule-dependent processes may have co-evolved. Consistent with this idea, closely related frog species inhabiting different temperature environments show distinct microtubule growth rates and catastrophe frequencies [31].

In summary, our comparison of *C. elegans* and *C. inopinata* demonstrates that modulation of microtubule regulatory mechanisms can affect nuclear and spindle dynamics during early embryogenesis. Further comparative analyses among closely related species will contribute to the understanding of the evolution of cellular dynamics.

## Acknowledgements

We thank members of the Sugimoto Lab for helpful discussion. Some strains were provided by the CGC, which is funded by NIH Office of Research Infrastructure Programs (P40 OD010440). This work was supported by Japan Science and Technology Agency (JST), Core Research for Evolutionary Science and Technology (CREST) Grant Number JPMJCR18S7 and Japan Society for the Promotion of Science (JSPS) KAKENHI Grant Number JP23K27152 to A.S., and JSPS KAKENHI Grant Number JP23KJ0148 and MEXT’s University Fellowship Founding Project for Innovation Creation in Science and Technology to S.O.

## Reference

1. Malone CJ, Misner L, Le Bot N, Tsai MC, Campbell JM, Ahringer J, et al. The C. elegans hook protein, ZYG-12, mediates the essential attachment between the centrosome and nucleus. Cell. 2003 Dec 26;115(7):825–36. doi: 10.1016/s0092-8674(03)00985-1. PMID: 14697201.

2. Kimura K, Kimura A. Intracellular organelles mediate cytoplasmic pulling force for centrosome centration in the Caenorhabditis elegans early embryo. Proc Natl Acad Sci U S A. 2011 Jan 4;108(1):137–42. doi: 10.1073/pnas.1013275108. Epub 2010 Dec 20. PMID: 21173218; PMCID: PMC3017145.

3. Park DH, Rose LS. Dynamic localization of LIN-5 and GPR-1/2 to cortical force generation domains during spindle positioning. Dev Biol. 2008 Mar 1;315(1):42–54. doi: 10.1016/j.ydbio.2007.11.037. Epub 2007 Dec 14. PMID: 18234174; PMCID: PMC2372164.

4. Grill SW, Gönczy P, Stelzer EH, Hyman AA. Polarity controls forces governing asymmetric spindle positioning in the Caenorhabditis elegans embryo. Nature. 2001 Feb 1;409(6820):630–3. doi: 10.1038/35054572. PMID: 11214323.

5. Tsou MF, Hayashi A, DeBella LR, McGrath G, Rose LS. LET-99 determines spindle position and is asymmetrically enriched in response to PAR polarity cues in C. elegans embryos. Development. 2002 Oct;129(19):4469–81. doi: 10.1242/dev.129.19.4469. PMID: 12223405.

6. Labbé JC, McCarthy EK, Goldstein B. The forces that position a mitotic spindle asymmetrically are tethered until after the time of spindle assembly. J Cell Biol. 2004 Oct 25;167(2):245–56. doi: 10.1083/jcb.200406008. Epub 2004 Oct 18. PMID: 15492042; PMCID: PMC2172534.

7. Gusnowski EM, Srayko M. Visualization of dynein-dependent microtubule gliding at the cell cortex: implications for spindle positioning. J Cell Biol. 2011 Aug 8;194(3):377–86. doi: 10.1083/jcb.201103128. PMID: 21825072; PMCID: PMC3153651.

8. Pecreaux J, Röper JC, Kruse K, Jülicher F, Hyman AA, Grill SW, Howard J. Spindle oscillations during asymmetric cell division require a threshold number of active cortical force generators.Curr Biol. 2006 Nov 7;16(21):2111–22. doi: 10.1016/j.cub.2006.09.030. PMID: 17084695.

9. Goldstein B. On the evolution of early development in the Nematoda. Philos Trans R Soc Lond B Biol Sci. 2001 Oct 29;356(1414):1521–31. doi: 10.1098/rstb.2001.0977. PMID: 11604120; PMCID: PMC1088533.

10. Brauchle M, Kiontke K, MacMenamin P, Fitch DH, Piano F. Evolution of early embryogenesis in rhabditid nematodes. Dev Biol. 2009 Nov 1;335(1):253–62. doi: 10.1016/j.ydbio.2009.07.033. Epub 2009 Jul 28. PMID: 19643102; PMCID: PMC2763944.

11. Valfort AC, Launay C, Sémon M, Delattre M. Evolution of mitotic spindle behavior during the first asymmetric embryonic division of nematodes. PLoS Biol. 2018 Jan 22;16(1):e2005099. doi: 10.1371/journal.pbio.2005099. PMID: 29357348; PMCID: PMC5794175.

12. Kanzaki N, Tsai IJ, Tanaka R, Hunt VL, Liu D, Tsuyama K, et al. Biology and genome of a newly discovered sibling species of Caenorhabditis elegans. Nat Commun. 2018 Aug 10;9(1):3216. doi: 10.1038/s41467-018-05712-5. PMID: 30097582; PMCID: PMC6086898.

13. Brenner S. The genetics of Caenorhabditis elegans. Genetics. 1974 May;77(1):71–94. doi: 10.1093/genetics/77.1.71. PMID: 4366476; PMCID: PMC1213120.

14. Oomura S, Tsuyama K, Haruta N, Sugimoto A. Transgenesis of the gonochoristic nematode Caenorhabditis inopinata by microparticle bombardment with hygromycin B selection. MicroPubl Biol. 2022 May 5;2022: 10.17912/micropub.biology.000564. doi: 10.17912/micropub.biology.000564. PMID: 35622530; PMCID: PMC9073556.

15. Toya M, Iida Y, Sugimoto A. Imaging of mitotic spindle dynamics in Caenorhabditis elegans embryos. Methods Cell Biol. 2010;97:359–72. doi: 10.1016/S0091-679X(10)97019-2. PMID: 20719280.

16. Altschul SF, Gish W, Miller W, Myers EW, Lipman DJ. Basic local alignment search tool. J Mol Biol. 1990 Oct 5;215(3):403–10. doi: 10.1016/S0022-2836(05)80360-2. PMID: 2231712.

17. Altschul SF,Madden TL, Schäffer AA, Zhang J, Zhang Z, Miller W, et al. Gapped BLAST and PSI-BLAST: a new generation of protein database search programs. Nucleic Acids Res. 1997 Sep 1;25(17):3389–402. doi: 10.1093/nar/25.17.3389. PMID: 9254694; PMCID: PMC146917.

18. Schäffer AA, Aravind L, Madden TL, Shavirin S, Spouge JL, Wolf YI, et al. Improving the accuracy of PSI-BLAST protein database searches with composition-based statistics and other refinements. Nucleic Acids Res. 2001 Jul 15;29(14):2994–3005. doi: 10.1093/nar/29.14.2994. PMID: 11452024; PMCID: PMC55814.

19. Maeda I, Kohara Y, Yamamoto M, Sugimoto A. Large-scale analysis of gene function in Caenorhabditis elegans by high-throughput RNAi. Curr Biol. 2001 Feb 6;11(3):171–6. doi: 10.1016/s0960-9822(01)00052-5. PMID: 11231151.

20. Kamath RS, Martinez-Campos M, Zipperlen P, Fraser AG, Ahringer J. Effectiveness of specific RNA-mediated interference through ingested double-stranded RNA in Caenorhabditis elegans. Genome Biol. 2001;2(1):RESEARCH0002. doi: 10.1186/gb-2000-2-1-research0002. Epub 2000 Dec 20. PMID: 11178279; PMCID: PMC17598.

21. Sturm Á, Saskoi É, Tibor K, Weinhardt N, Vellai T. Highly efficient RNAi and Cas9-based auto-cloning systems for C. elegans research. Nucleic Acids Res. 2018 Sep 28;46(17):e105. doi: 10.1093/nar/gky516. PMID: 29924347; PMCID: PMC6158509.

22. Riche S, Zouak M, Argoul F, Arneodo A, Pecreaux J, Delattre M. Evolutionary comparisons reveal a positional switch for spindle pole oscillations in Caenorhabditis embryos. J Cell Biol. 2013 May 27;201(5):653–62. doi: 10.1083/jcb.201210110. Epub 2013 May 20. PMID: 23690175; PMCID: PMC3664713.

23. Enos SJ, Dressler M, Gomes BF, Hyman AA, Woodruff JB. Phosphatase PP2A and microtubule-mediated pulling forces disassemble centrosomes during mitotic exit. Biol Open. 2018 Jan 12;7(1):bio029777. doi: 10.1242/bio.029777. PMID: 29222174; PMCID: PMC5829501.

24. Magescas J, Zonka JC, Feldman JL. A two-step mechanism for the inactivation of microtubule organizing center function at the centrosome. Elife. 2019 Jun 27;8:e47867. doi: 10.7554/eLife.47867. PMID: 31246171; PMCID: PMC6684319.

25. O’Rourke SM, Christensen SN, Bowerman B. Caenorhabditis elegans EFA-6 limits microtubule growth at the cell cortex. Nat Cell Biol. 2010 Dec;12(12):1235–41. doi: 10.1038/ncb2128. Epub 2010 Nov 14. PMID: 21076413; PMCID: PMC3236679.

26. Cluet D, Stébé PN, Riche S, Spichty M, Delattre M. Automated high-throughput quantification of mitotic spindle positioning from DIC movies of Caenorhabditis embryos. PLoS One. 2014 Apr 24;9(4):e93718. doi: 10.1371/journal.pone.0093718. PMID: 24763198; PMCID: PMC3998942.

27. Han X, Adames K, Sykes EM, Srayko M. The KLP-7 Residue S546 Is a Putative Aurora Kinase Site Required for Microtubule Regulation at the Centrosome in C. elegans. PLoS One. 2015 Jul 13;10(7):e0132593. doi: 10.1371/journal.pone.0132593. PMID: 26168236; PMCID: PMC4500558.

28. Connolly AA, Sugioka K, Chuang CH, Lowry JB, Bowerman B. KLP-7 acts through the Ndc80 complex to limit pole number in C. elegans oocyte meiotic spindle assembly. J Cell Biol. 2015 Sep 14;210(6):917–32. doi: 10.1083/jcb.201412010. PMID: 26370499; PMCID: PMC4576866.

29. Hatanaka R, Tamagawa K, Haruta N, Sugimoto A. The impact of differential transposition activities of autonomous and nonautonomous hAT transposable elements on genome architecture and gene expression in Caenorhabditis inopinata. Genetics. 2024 Jun 5;227(2):iyae052. doi: 10.1093/genetics/iyae052. PMID: 38577765; PMCID: PMC11492494.

30. Kawahara K, Inada T, Tanaka R, Dayi M, Makino T, Maruyama S, et al. Differentially Expressed Genes Associated with Body Size Changes and Transposable Element Insertions between Caenorhabditis elegans and Its Sister Species, Caenorhabditis inopinata. Genome Biol Evol. 2023 Apr 6;15(4):evad063. doi: 10.1093/gbe/evad063. PMID: 37071793; PMCID: PMC10139442.

31. Hirst WG, Biswas A, Mahalingan KK, Reber S. Differences in Intrinsic Tubulin Dynamic Properties Contribute to Spindle Length Control in Xenopus Species. Curr Biol. 2020 Jun 8;30(11):2184–2190.e5. doi: 10.1016/j.cub.2020.03.067. Epub 2020 May 7. PMID: 32386526.

